# Scale Reliant Mixed Effects Models Enhance Microbiome Data Analysis

**DOI:** 10.1101/2025.08.05.668734

**Authors:** Kyle C. McGovern, Justin D. Silverman

## Abstract

Linear models, including those used for differential abundance analyses, are frequently used in microbiome research to assess how experimental conditions (e.g., disease state or age) affect microbial abundance. Linear mixed-effects models (MEMs) extend linear models to accommodate complex designs, such as longitudinal sampling or hierarchical study structures. However, when applied to microbiome data, existing MEM approaches suffer from high false positive and false negative rates because sequence counts are compositional – they reflect relative rather than absolute abundances. Current methods attempt to overcome this limitation through normalization, but these approaches imply on strong, often unrealistic assumptions about the unmeasured biological scale (e.g., total microbial load). Here we introduce scale-reliant mixed-effects models (SR-MEM), which extend our earlier scale-reliant inference framework by explicitly modeling uncertainty in the unmeasured scale via user-defined probability distributions. By treating scale as a latent variable rather than fixing it through normalization, SR-MEM yields robust inference for complex experimental designs. Across simulations and real datasets, SR-MEM is the only method that consistently controls the false discovery rate while achieving higher power than both normalization and bias-correction methods. SR-MEM can also incorporate external scale measurements (e.g., flow cytometry, qPCR) or leverage independent studies to further improve inference. An accessible implementation is provided in the ALDEx3 R package, enabling more rigorous and reproducible analysis of microbial communities.

## 1 Background

A key goal in microbiome data analysis (e.g., 16S rRNA-seq or shotgun metagenomics) is to identify how experimental and biological factors (e.g., health vs. disease) affect the abundance of different taxa. Linear models are widely used for this task. Yet current linear modeling methods often inflate false-positive and false-negative rates because they cannot accommodate complex study designs or account for limitations of the sequencing measurement process [1, 2, 3].

Linear mixed-effects models (MEMs) extend linear models by incorporating random effects to model structured sources of variation, improving inference in hierarchical or longitudinal studies. For example, random intercepts can account for cage effects in co-housed mice [4, 5], batch effects and contamination [6, 7, 8], or temporal autocorrelation in repeated measurements [9, 10]. Several MEM-based tools have been developed for sequence count data, including NBZIMM [11], MaAsLin2 [12], lmerSeq [13], ANCOM-BC2 [14], and LinDA [15].

However, these methods remain fundamentally limited by compositionality: sequencing data capture only relative–not absolute–abundances and provide no information about the total biological scale (e.g., microbial load), which is essential for accurate inference [3, 16, 17]. Most methods attempt to circumvent this with normalizations such as Total Sum Scaling (MaAsLin2), mean-variance weighting (lmerSeq, limma-voom), or sequencing-depth offsets (NBZIMM). Yet normalizations impose strong, hidden assumption about scale, biasing estimates and inflating error rates [2, 18, 19]. More recent bias-correction approaches, such as ANCOM-BC2 and LinDA, still depend on unverifiable assumptions (e.g., that a stable set of taxa remains unchanged in absolute abundance across conditions) and fail when those assumptions are violated [15, 20].

Here, we introduce Scale-Reliant Mixed-Effects Models (SR-MEM), a general framework for mixed-effects modeling of microbiome sequence count data. SR-MEM extends our recently developed Scale-Reliant Inference (SRI) framework, which uses specialized Bayesian partially identified models (PIMs) to avoid normalization and instead explicitly model uncertainty in the unmeasured biological scale through a user-defined prior, termed a scale model. By treating scale as an uncertain latent variable rather than assuming it can be recovered through normalization, SR-MEM provides a flexible and robust framework for mixed-effects modeling that is robust to errors in scale assumptions. Across simulations and real data analyses, SR-MEM consistently reduces false-positive and false-negative rates compared to existing methods. For example, in reanalyses of a longitudinal study on antibiotic effects in the gut microbiome, SR-MEM corrected previous failures in which normalization-based methods erroneously concluded that carbapenems had no effect other than increasing a single taxon – despite this class of antibiotics having broad activity against gut bacteria. Moreover, SR-MEM improves reproducibility: in replication studies, SR-MEM produces results consistent with prior findings, whereas normalization-based methods yield contradictory conclusions. We provide an open-source implementation of SR-MEM in the ALDEx3 R package, making the method broadly accessible for rigorous and reproducible mixed effects modeling of microbiome data.

## 2 Results

### 2.1 Scale-Reliant Mixed-Effects Models (SR-MEM)

SR-MEM builds on the theory of *Scale-Reliant Inference (SRI)*, which provides a principled framework for estimating quantities that depend on absolute abundances when only noisy, compositional count data are observed [3].

#### 2.1.1 The SRI Framework

Let *W* denote a *D × N* matrix of absolute abundances, where *W*_*dn*_ is the abundance of taxon *d* in biological system *n* (e.g., a gut microbiome). *W* can be uniquely described by its composition (relative abundances) and scale (total abundances) via:

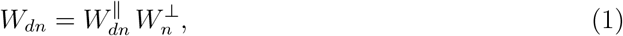

where *W*^∥^ is the composition: a *D × N* matrix of proportions satisfying 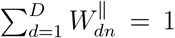 for each *n*; and *W* ^*⊥*^ is the scale: a positive-valued *N*-vector of total abundances satisfying 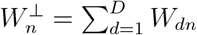.

In practice, *W* is not directly observed. Instead, we observe a *D×N* count table *Y*, where *Y*_*dn*_ is the number of sequencing reads assigned to taxon *d* in sample *n*. These counts provide noisy, sparse information about the composition *W*^∥^ but little to no information about the scale *W* ^*⊥*^. More formally, SRI defines “little to no information” in terms of identification restrictions [3].

The goal of an SRI analysis is formalized as a *target estimand*: a quantity *θ*:= *θ*(*W*) that depends on the unobserved absolute abundances *W* and may therefore depend on both the composition *W*^∥^ and the scale *W* ^*⊥*^. Prior work has studied a range of estimands under the SRI framework, including the gene set enrichment estimand [19], correlations between taxa [3], and log-fold changes [2, 3, 19]. Notably, log-fold changes are themselves a special case of linear model estimands, which in turn are special cases of the mixed-effects model estimands we introduce next.

#### 2.1.2 The Target Estimand in SR-MEM

SR-MEM extends SRI to mixed-effects modeling, enabling inference under complex experimental designs. Suppose we had access to the absolute abundances *W*. For each taxon *d*, our goal would be to estimate the fixed-effect parameters *θ*_*d*_ in the standard linear mixed-effects model:

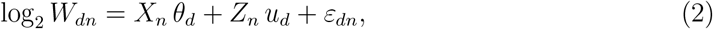

where:

- *X*_*n*_ is a *P* -vector of fixed-effect covariates (observed),
- *θ*_*d*_ is the *P* -vector of *fixed effects* (the target estimand; estimated),
- *Z*_*n*_ is a *Q*-vector of random-effect covariates (observed; e.g., subject, cage, collection site),
- *u*_*d*_ *∼ 𝒩* (0, *G*) captures structured random variation (estimated), and
- *ε*_*dn*_ *∼ 𝒩* (0, *R*_*ρ*_) is a residual error term (estimated).

More formally, the target estimand is actually an estimator of *θ*_*d*_ within this model (see *Methods*). We model *W* on the log_2_ scale because absolute abundances are non-negative and microbial communities are more naturally described on a log scale, where changes are often multiplicative rather than additive [21].

A full review of linear mixed-effects models is beyond the scope of this work (see, [22] for a general overview and [23] for a narrower overview with application to microbiome data analysis). Here, we briefly illustrate how the interpretation of the fixed effects *θ*_*d*_ depends on the choice of fixed-effect covariates *X*_*n*_, random-effect covariates *Z*_*n*_, and the covariance structures *G* and *R*_*ρ*_, all of which are determined by the experimental design:

- **Binary covariates**. If *X*_*n*_ = (1, *x*_*n*_), where *x*_*n*_ = 0 for control and *x*_*n*_ = 1 for treatment, then *θ*_*d*_ = (*α*_*d*_, *β*_*d*_), with *α*_*d*_ the baseline log_2_-abundance in the control group and *β*_*d*_ the log_2_-fold change in absolute abundance between treatment and control. That is, *β*_*d*_ is the standard estimand used in differential abundance analyses.
- **Continuous covariates**. If *X*_*n*_ = (1, age_*n*_), then *θ*_*d*_ = (*α*_*d*_, *γ*_*d*,age_), where *γ*_*d*,age_ represents the change in log_2_-abundance per unit increase in age, holding other covariates constant.
- **Longitudinal studies**. If *X*_*n*_ = (1, treatment_*n*_), where treatment_*n*_ = 0 for control and 1 for treatment, then *θ*_*d*,treatment_ represents the average log_2_-fold change in absolute abundance between groups. Suppose each subject is measured repeatedly over time. Then *R*_*ρ*_ is often specified with an AR1 structure, where for two samples *n* and 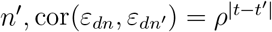, to account for temporal correlations within subjects.
- **Hierarchical or nested designs**. In clustered designs (e.g., mice co-housed in cages), *Z*_*n*_ could be a vector of cluster-level indicators (e.g., cage membership), and *u*_*d*_ could represent the corresponding random effects. Together, *Z*_*n*_*u*_*d*_ accounts for variation introduced by the hierarchy, such as cage-to-cage differences. The fixed effects *θ*_*d*_ are then interpreted as population-level effects after adjusting for this structured variability.

### 2.2 Inference in SR-MEM

Our goal is to estimate the fixed effects *θ*_*d*_ from Eq. (2) as if the absolute abundances *W* were observed. Two sources of uncertainty complicate this task. First, the observed sequence counts *Y* are noisy and sparse, introducing uncertainty in the compositional component *W*^∥^. Second, and more fundamentally, the scale component *W* ^*⊥*^ is unidentifiable from *Y* alone because sequence counts provide no direct information about total microbial load.

To address the first source of uncertainty, we follow the strategy used in ALDEx2: *N* independent Bayesian multinomial-Dirichlet models are fit to the observed counts *Y*, and posterior samples from these models are treated as plausible realizations of the underlying composition *W*^∥^ (see *Methods*). The key innovation of SRI, and by extension SR-MEM, is to also account for uncertainty in the unobserved scale *W* ^*⊥*^—a source of variation that normalization-based methods typically ignore. Standard normalizations implicitly assume that *W* ^*⊥*^ can be recovered without error (e.g., through sequencing depth offsets or total-sum scaling) [2, 3, 18]. Because *Y* contains no scale information, any single imputation of *W* ^*⊥*^ introduces bias. Instead, SRI introduces a *scale model P*: a user-defined probability distribution over *W* ^*⊥*^ that explicitly encodes prior knowledge and uncertainty about scale. This formulation places SR-MEM within the broader class of *Bayesian partially identified models* (PIMs), where inference proceeds by integrating over distributions of unidentifiable components rather than fixing them [3].

SR-MEM inference proceeds by sampling separate realizations of *W*^∥^ (from multinomial-Dirichlet models) and *W* ^*⊥*^ (from the scale model *P*). These are combined via Eq. (1) to generate realizations of the absolute abundances *W*. For each realization of *W*, a mixed-effects model (Eq. 2) is fit, yielding a corresponding realization of the target estimand *θ*_*d*_. Repeating this procedure across Monte Carlo samples produces a posterior distribution for *θ*_*d*_ and for the corresponding *p*-values (adjusted for multiple testing as described in *Methods*).

Overall the SR-MEM inference procedure is intuitive, combining uncertainty about *W*^∥^ and *W* ^*⊥*^ during inference of *θ*_*d*_. This inference procedure is also theoretically rigorous. Together this procedure defines a Scale Simulation Random Variable (SSRV) which is itself an example of a Bayesian Partially Identified Model [3].

### 2.3 Designing Scale Models for SR-MEM

At first glance, defining a scale model for 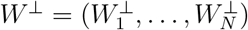 may seem daunting because it appears to require specifying a joint distribution over *N* unobserved scale parameters. However, the field of Scale-Reliant Inference (SRI) has developed practical strategies to make this task tractable. In Supplementary File 1, we review three general approaches: (1) constructing scale models directly from external measurements (e.g., flow cytometry or qPCR); (2) reformulating standard normalization procedures as scale models; and (3) avoiding explicit modeling of the full vector *W* ^*⊥*^ by instead specifying how covariates affect scale.

To illustrate the third approach, consider a longitudinal study with repeated, biweekly measurements of the oral microbiome before and after tooth brushing. The goal is to identify taxa that change in abundance after brushing (differential abundance analysis). Rather than specifying a full joint distribution over *W* ^*⊥*^, we can define a simpler model for the change in scale between conditions. Let

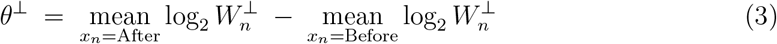

denote the difference in average log-scale between the two conditions. Because oral microbial load is expected to decrease after brushing, we anticipate *θ*^*⊥*^ *<* 0. A simple choice is an asymmetric prior,

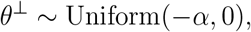

with *α* chosen conservatively based on prior literature. For instance, brushing has been reported to reduce oral microbial load by up to 99%, corresponding to *α* = 6.6 [24, 25]. As discussed in *Methods*, specifying a model for *θ*^*⊥*^ implicitly defines a model for *W* ^*⊥*^ through its relationship in Equation (3). Thus we have simplified the problem: we implicitly specify a scale model over 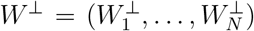 by specifying a simpler, one-dimensional, model for *θ*^*⊥*^.

### 2.4 SR-MEM Substantially Reduces False Positives while Maintaining Power

We benchmarked SR-MEM against five widely used mixed-effects modeling tools: three employing distinct normalization strategies (NBZIMM [11], MaAsLin2 [12], and lmerSeq [13]) and two using bias-correction approaches (ANCOM-BC2 [14] and LinDA [15]). Simulated 16S rRNA count data were generated with SparseDOSSA2, which reproduces realistic sparsity, variance, and taxon–taxon correlations based on real microbiome data [26]. We selected SparseDOSSA2 because its generative model is independent of all methods tested, including SR-MEM. Two simulation models were used, trained on oral [27] and skin [28] microbiome datasets. To make the scenarios biologically interpretable, we modeled a hypothetical pre-biotic intervention in which participants in the treatment arm received a broad-spectrum prebiotic expected to increase total microbial load in the gut.

We simulated studies with two equally sized groups (treatment and control), 6–300 participants, and 20 repeated measurements per participant (see *Methods*). Three scenarios were evaluated: two based on the oral microbiome model with 50% and 80% of taxa differentially abundant (DA) and one based on the skin microbiome model with 50% DA. The oral microbiome simulations included on average *≈* 135 taxa, whereas the skin microbiome simulation included *≈* 35. The number of taxa varied slightly between simulations due to sparsity-based filtering, which removed different taxa in each simulation (see *Methods*). Across all scenarios, the average log-scale difference in total microbial load between conditions was within *θ*^*⊥*^ *∈* [1, 2]. For SR-MEM, we used a moderately informative scale model *θ*^*⊥*^ *∼ 𝒩* (1.3, 0.5^2^), which reflected an assumption that, with 95% probability *θ*^*⊥*^ *∈* [0.32, 2.27].

SR-MEM was the only method that consistently controlled FDR across all study sizes and scenarios (Fig. 1). Supplementary Figure 1 compares SR-MEM to ALDEx3 using the same scale model but with linear modeling (as implemented in ALDEx2). Unlike SR-MEM, ALDEx3 with linear modeling failed to control FDR, demonstrating that SR-MEM’s performance stems from integrating the scale model with mixed-effects modeling.

**Figure 1:**
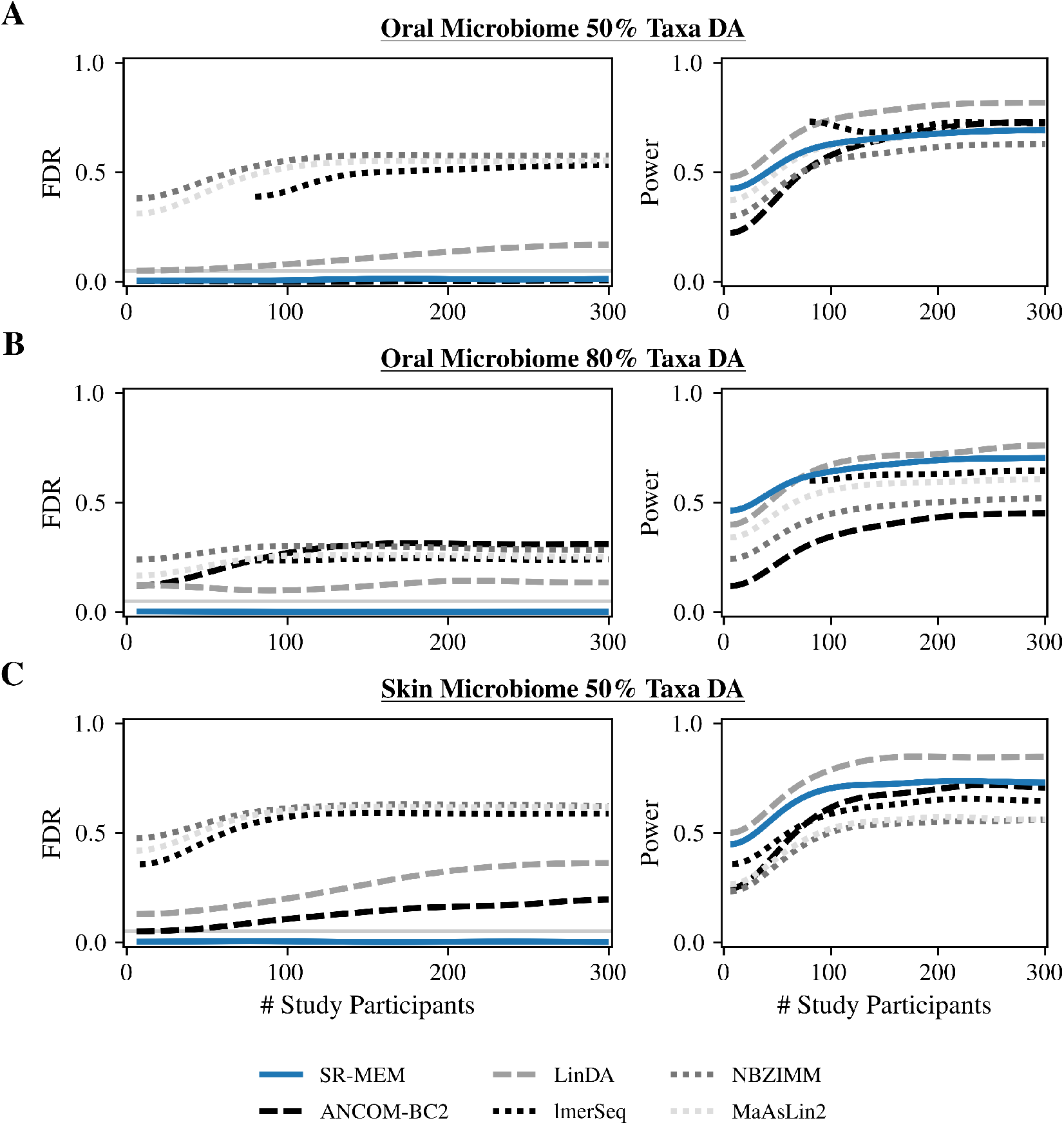
SR-MEM uniquely controls false discoveries across diverse study designs while maintaining power. We benchmarked SR-MEM (solid blue line) against three normalization-based methods (lmerSeq, NBZIMM, MaAsLin2; black/gray dotted lines) and two bias-correction methods (ANCOM-BC2, LinDA; black/gray dashed lines). Simulated data were generated with SparseDOSSA2 trained on oral and skin microbiome datasets [27, 28]. False discovery rate (FDR) and power were averaged over 20 simulations. Only SR-MEM consistently maintained FDR at or below the nominal 0.05 level (gray horizontal line). All methods used the Benjamini–Hochberg procedure for multiple hypothesis correction. **A**. Oral microbiome model with 50% DA taxa. **B**. Oral microbiome model with 80% DA taxa. **C**. Skin microbiome model with 50% DA taxa.

Bias-correction methods (ANCOM-BC2, LinDA) failed to consistently control FDR. ANCOM-BC2 pools information across taxa to estimate scale differences between conditions, whereas LinDA estimates bias from centered log-ratio (CLR) transformed data using the mode of estimated fixed effects. Both approaches failed to reliably recover bias terms from observed counts. ANCOM-BC2 controlled FDR only in the oral microbiome scenario with 50% DA taxa (Fig. 1A), likely because a larger proportion of non-DA taxa stabilized its bias estimation. Its performance deteriorated when most taxa were DA (Fig. 1B) or when fewer taxa were present (Fig. 1C). SR-MEM generally had higher power than ANCOM-BC2, and although LinDA occasionally achieved slightly higher power, it consistently failed to control FDR. In smaller studies (*<* 80 participants) with 80% DA taxa, SR-MEM outperformed LinDA in both FDR and power (Fig. 1B).

Normalization-based methods consistently exhibited inflated FDR and generally lower power. NBZIMM’s failure aligns with prior findings that zero-inflation assumptions can increase false positives and negatives [29]. MaAsLin2 uses total-sum scaling, implicitly assuming no change in microbial load between conditions (*θ*^*⊥*^ = 0), while lmerSeq normalizes to a pseudo-reference derived from observed counts before applying a variance-stabilizing transformation [30, 11]. These assumptions introduce bias, leading to inflated false discoveries.

Overall, SR-MEM was the only method to consistently maintain FDR control while achieving competitive or superior power across realistic microbiome study designs.

### 2.5 SR-MEM Enhances Statistical Power in an Oral Microbiome Study

To assess SR-MEM’s performance on real data, we reanalyzed an oral microbiome dataset where unstimulated saliva samples were collected from 28 participants across four perturbations: water (control), antiseptic mouthwash, alcohol-free mouthwash, and soda [31]. Microbial composition and total microbial load (scale) were measured via 16S rRNA sequencing and flow cytometry at three time points: baseline, 15 minutes post-perturbation, and two hours post-perturbation. Two technical replicate flow cytometry measurements of total salivary microbial load were taken for each sample. We used these flow cytometry measurements to define a scale model that accounted for the technical variability in the measurement process (see *Methods*). Using SR-MEM, we specified a mixed effects model with time, perturbation, and their interaction as fixed effects and with a participant-specific random intercept. Figure 2 shows results for the effect of alcohol-free mouthwash 15 minutes post-perturbation, the only condition with significant genus-level absolute abundance changes. See Supplementary File 2 for full results.

**Figure 2:**
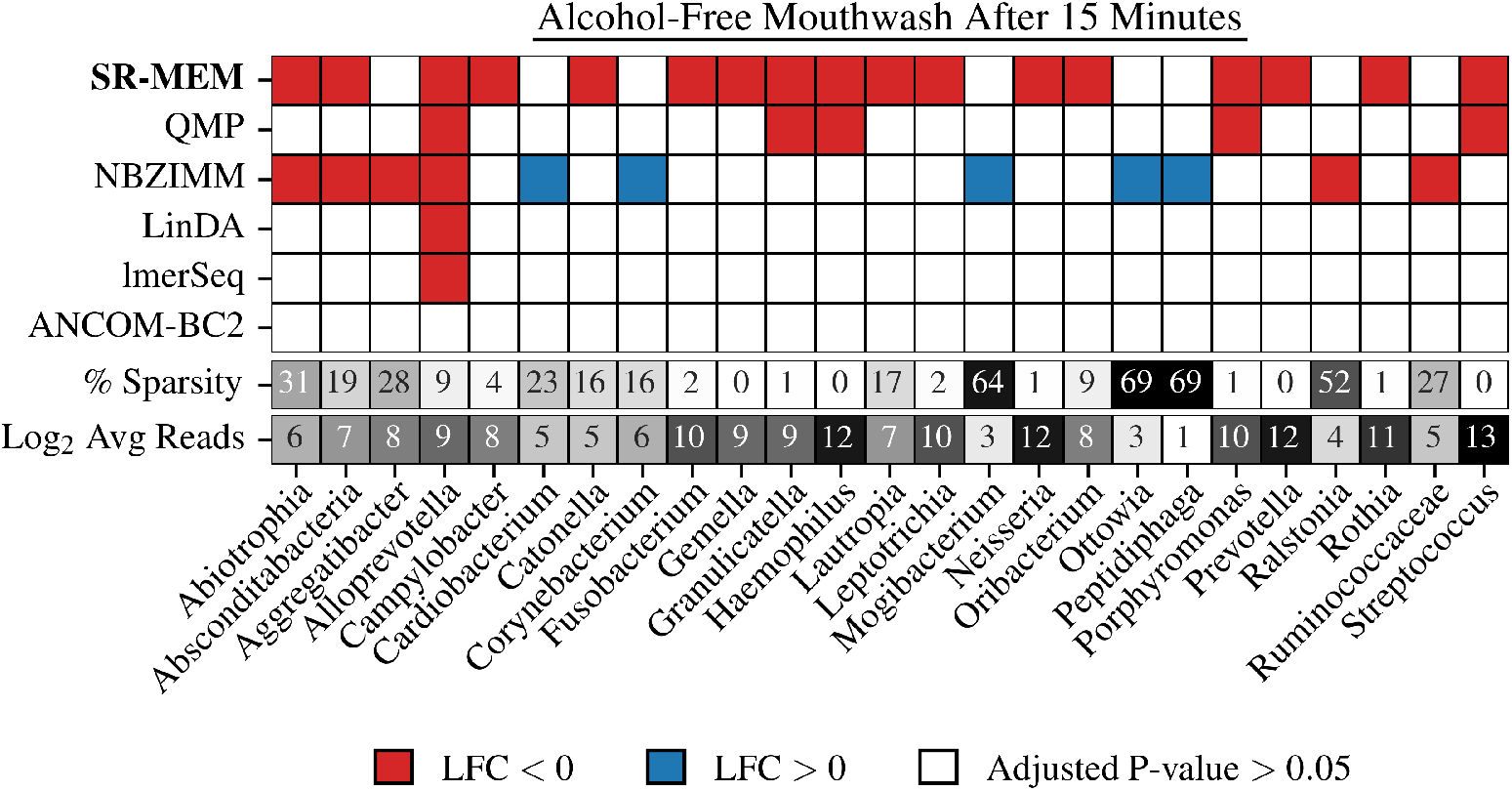
SR-MEM improves detection of genera that differ in abundance after treatment with alcohol-free mouthwash. SR-MEM and QMP leveraged external microbial load measurements, while NBZIMM, LinDA, lmerSeq, and ANCOM-BC2 relied on normalization and bias correction. All methods used a mixed-effects model with perturbation, time point, and their interaction as fixed effects and a participant-specific random intercept. *p*-values were adjusted for multiple hypothesis testing using the Benjamini-Hochberg procedure. Sparsity indicates the percentage of samples where a given taxon was unobserved (zero counts), the last row represents the log_2_-transformed mean read count per taxon across all samples.

SR-MEM identified 17 genera which decreased in abundance 15 minutes after alcohol-free mouthwash (Fig. 2). Decreasing microbial abundances after mouthwash is both intuitive and aligns with flow cytometry data which suggested a 3.8-log decrease in microbial load in the post-mouthwash condition. We compared the SR-MEM results to four alternative methods: Quantitative Microbial Profiling (QMP), which also used flow cytometry data, and four normalization- and bias correction-based approaches: ANCOM-BC2, LinDA, lmerSeq, and NBZIMM [16].

QMP identified only 5 of the 17 genera detected by SR-MEM. The reduced power of QMP compared to SRI-based methods is well established [3] and likely stems from differences in how the two approaches model the compositional component *W*^∥^. Both QMP and SR-MEM incorporate flow cytometry data to inform scale modeling; however, QMP first rarefies the data and then applies total-sum scaling to estimate *W*^∥^. Rarefaction discards a substantial fraction of reads, decreasing statistical power [32, 3], whereas SR-MEM uses a Bayesian multinomial–Dirichlet model that retains all available counts and has been shown to improve power [3]. Consistent with this expectation, rarefaction in QMP increased the sparsity of the count matrix *Y* from 37% to 69%, indicating that a large proportion of data was effectively discarded.

The normalization- and bias-correction methods produced results consistent with their simulated performance (Fig. 1). ANCOM-BC2 showed lower power than SR-MEM and detected no significant genus-level changes. LinDA and lmerSeq each identified only *Alloprevotella* as decreasing. In contrast, NBZIMM frequently flagged high-sparsity, low-count genera as differentially abundant, likely inflating false positives (Fig. 2). For instance, it reported increases in *Mogibacterium, Ottowia*, and *Peptidiphaga*, taxa with 64–69% sparsity, contradicting both the expected antimicrobial effects of alcohol-free mouthwash and the observed 3.8-log decrease in total microbial load [33]. These findings mirror our simulation results, where NBZIMM consistently exhibited the highest FDR.

Together, these results indicate that in real data, SR-MEM improves power compared to alternative methods and can decrease false discovery rates.

### 2.6 SR-MEM Yields More Reproducible Results Than Normalization-Based Methods

To evaluate the reproducibility of SR-MEM, we compared its results on a large metagenomic cohort of Crohn’s disease patients (the Inflammatory Bowel Disease Multi-omics Database; IBDMDB) to results from a smaller, independent 16S rRNA study (Vandeputte et al. [16]). The IBDMDB dataset comprised 944 samples from 27 healthy and 50 Crohn’s disease (CD) subjects, whereas the Vandeputte study included 95 samples from 29 healthy and 66 CD subjects. The Vandeputte study, which included paired flow cytometry measurements, was previously analyzed using scale reliant linear models [2]—appropriate for its cross-sectional design with a single measurement per subject. In contrast, the IBDMDB data include repeated biweekly measurements collected over one year across five body sites, making those earlier linear-model-based SRI methods inappropriate. Instead, mixed-effects modeling is required to account for participant- and site-specific variation. We therefore treated the Vandeputte results as a reference and evaluated the reproducibility of SR-MEM in this more complex longitudinal metagenomic setting.

Because the IBDMDB lacked paired flow cytometry or qPCR measurements, we followed Nixon et al. [2] and defined a scale model based on prior literature which specifically studied how total microbial load varied between healthy subjects and subjects with Crohn’s disease [34] (see *Methods*). This scale model was centered at 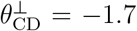, suggesting total microbial load is decreased in Crohn’s disease and has been independently validated [2]. The IBDMDB data included biweekly samples collected for one year across five body sites, with each sample classified as dysbiotic or non-dysbiotic to reflect the episodic nature of CD [35]. We applied SR-MEM with fixed effects for antibiotic use and a disease–dysbiosis interaction, including participant- and site-specific random intercepts. Comparisons focused on dysbiotic samples from CD patients versus non-dysbiotic samples from healthy participants.

For comparison, we analyzed two alternative methods: (1) *ASR-MEM*, which used the same mixed-effects structure but applied arcsine square-root normalization following the original IBDMDB analysis [35]; and (2) *SR-MEM-CLR*, which used the same SR-MEM structure but replaced the literature-based scale model with Centered Log-Ratio (CLR) normalization as implemented in ALDEx2 [30]. The results are shown in Fig. 3.

**Figure 3:**
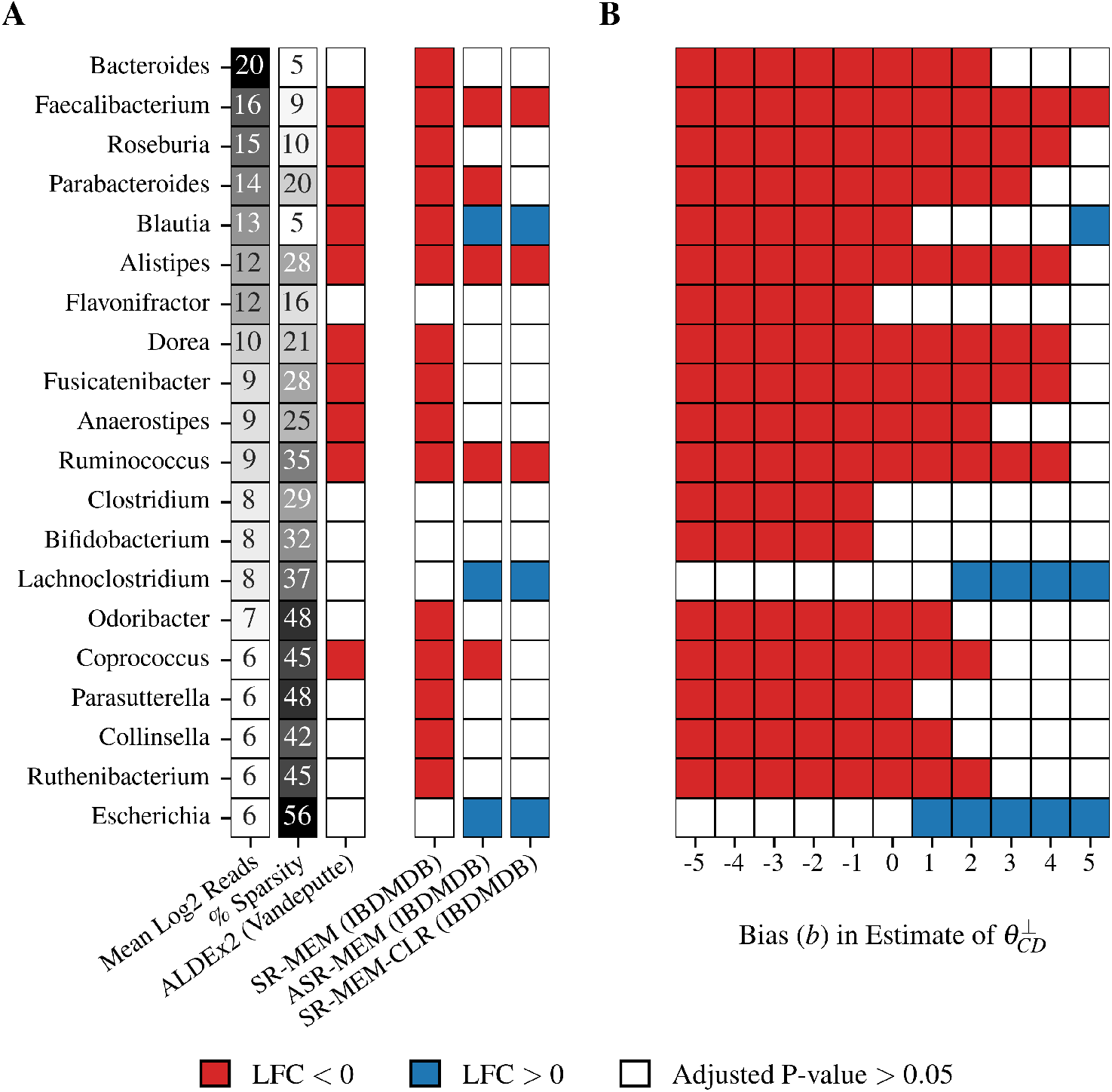
SR-MEM analysis of Crohn’s disease metagenomic data is more reproducible than normalization-based methods. **A**. Differential abundance results for the 25 genera with the highest mean log_2_ abundance (after adding a pseudo-count of 0.5 and applying a log_2_ transformation). SR-MEM, using a scale model based on microbial load estimates from Sarrabayrouse et al. [34], produced results consistent with a prior ALDEx2 analysis of the full Vandeputte dataset when applied to the IBDMDB cohort. In contrast, two alternative analyses of the same IBDMDB data—(1) *SR-MEM-CLR*, which replaced the scale model with Centered Log-Ratio (CLR) normalization, and (2) *ASR-MEM*, a mixed-effects model applied after arcsine square-root normalization as in the original IBDMDB study [35]—yielded results that differed both from SR-MEM and from the Vandeputte analysis. Notably, CLR normalization implicitly assumed a microbial load increase in CD patients *≈* of approximately 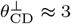, contradicting qPCR and flow cytometry measurements showing a substantial decrease [16, 34]. **B**. Sensitivity analysis of SR-MEM results. To assess the impact of potential misspecification of the scale model, a bias term *b* was introduced to shift the assumed scale difference to 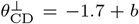, and differential abundance was re-estimated across *b ∈* [*−*5, 5]. The case *b* = 0 corresponds to the SR-MEM analysis in part A.

The SR-MEM analysis of the IBDMDB dataset produced results largely consistent with the prior Vandeputte analysis, albeit with higher power, likely due to the substantially larger IBDMDB sample size (Fig. 3A). Across all genera present in both studies, SR-MEM and the Vandeputte analysis agreed in both significance and direction of effect for all but five genera, which were significant in the IBDMDB analysis but not in Vandeputte. To assess whether these discrepancies reflected true positives (detected by SR-MEM due to increased power) or spurious false positives, we conducted a sensitivity analysis. Specifically, we introduced a bias parameter *b*, shifting the assumed scale difference to 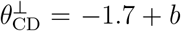 and re-estimated differential abundance across *b ∈* [*−*5, 5] (Fig. 3B). Two of the five genera, *Bacteroides* and *Ruthenibacterium*, were highly robust, remaining significant unless the scale model was misspecified by more than *b >* 2. Practically, *b* = 2 would imply an eight-fold higher microbial load in CD patients than supported by existing qPCR and flow cytometry data—an implausible scenario. These two genera therefore likely represent true positives that were undetected in the Vandeputte analysis due to either lower power or cohort-specific differences. In contrast, *Odoribacter, Collinsella*, and *Parasutterella* were more sensitive to perturbations in the scale model, suggesting that these results, while plausible, warrant further investigation and independent validation.

Whereas the results of SR-MEM were closely aligned to the prior analysis of Vandeputte, the results of ASR-MEM and SR-MEM-CLR were most similar to each other and markedly different than the Vandeputte analysis. Further investigation revealed that this similarity was primarily driven by normalization: in this study, CLR normalization implied the implausible assumption that gut microbial load was substantially increased in Crohn’s disease patients 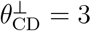, whereas qPCR and flow cytometry quantification of microbial load revealed a substantial decrease [16, 34]. This suggests that the normalizations used in SR-MEM-CLR and ASR-MEM likely introduced substantial bias. This conclusion is reinforced by noting that many of the differences between SR-MEM, ASR-MEM, and SR-MEM-CLR were highly sensitive to perturbations of the scale model (*b*). For example, ASR-MEM and SR-MEM-CLR suggest that *Blautia* is increased in Crohn’s but this result only holds in the unlikely scenario *b* = 5 [16, 34].

Our findings suggest that normalization-based methods can introduce substantial bias into analyses, degrading reproducibility. In contrast, SR-MEM combined with scale models built simply from prior literature can mitigate this bias and enhance reproducibility.

### 2.7 SR-MEM Provides More Accurate Antibiograms from Longitudinal Data

We reanalyzed a longitudinal study of 128 cancer patients hospitalized to receive allogeneic hematopoietic cell transplantation [36]. The dataset included 2,589 fecal samples with paired 16S rRNA sequencing and qPCR measurements, providing information on both gut microbiome composition and total microbial load. During hospitalization, patients received various oral and intravenous antibiotics. The original authors used these data to estimate antibi-ograms (tables summarizing the effect of each antibiotic on each gut microbe) but many of their findings were inconsistent with the known spectrum of activity of these drugs. We hypothesized that these inconsistencies arose from limitations in the statistical models used in the original analysis, and that applying SR-MEM would yield results more consistent with the known spectrum of antibiotic activity.

We used SR-MEM to estimate the fixed effect of each antibiotic. Subject-level random intercepts were used along with autocorrelated residuals to capture the temporal correlations between measurements taken from the same subject (see *Methods*). We used the paired qPCR measurements to define a scale model (see *Methods*). In Figure 4 we compare the results of SR-MEM with the results from the original authors methods for the antibiotics vancomycin and carbapenems [36]. A full comparison of all antibiotics are presented in Supplementary Figures 2 and 3.

**Figure 4:**
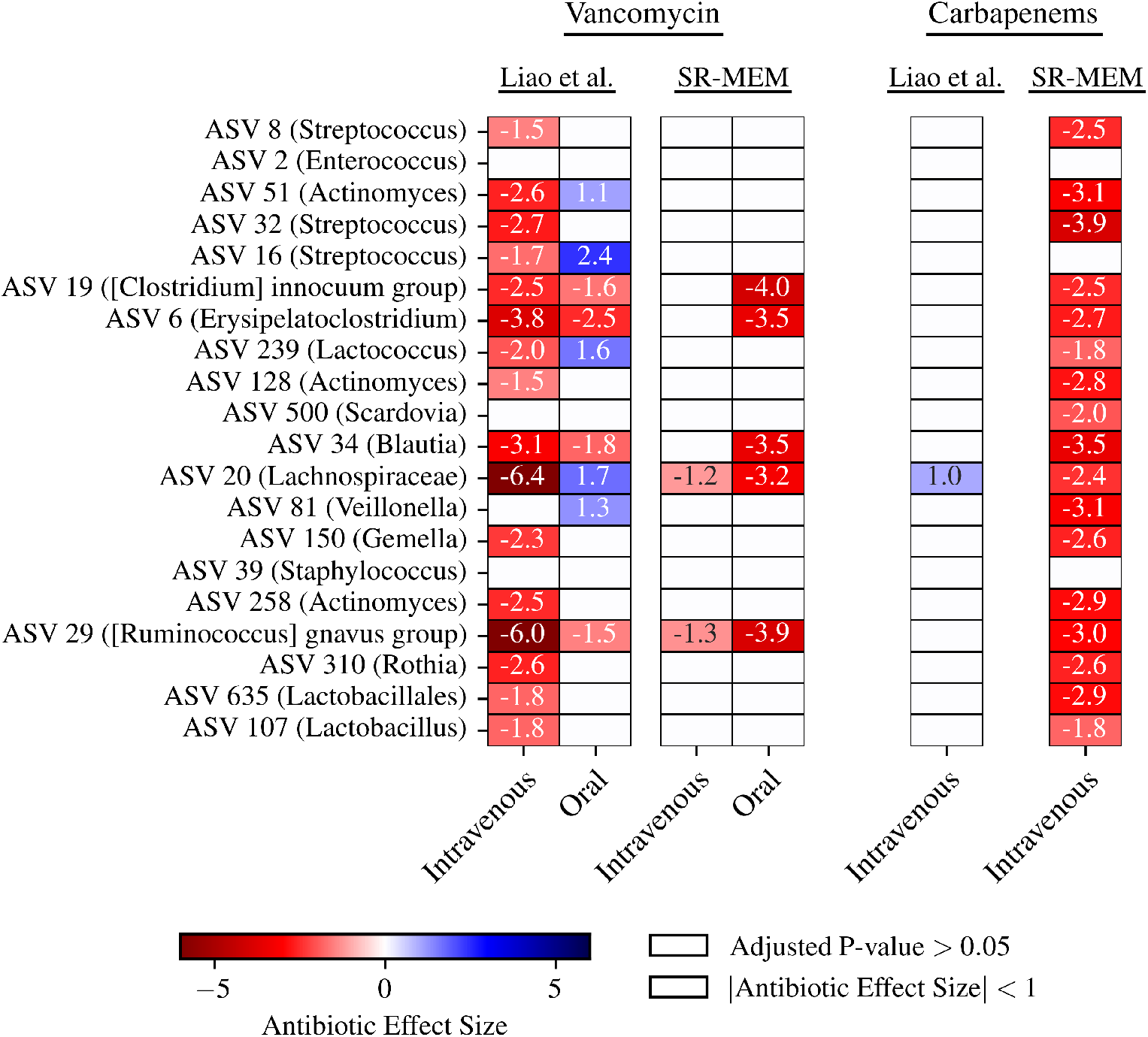
SR-MEM improves inference in a longitudinal study of antibiotic effects. Comparison of SR-MEM and the original analysis by Liao et al. [36] on the impact of orally- and intravenously-administered vancomycin and intravenously-administered carbapenems on the 20 most abundant gut microbial taxa at the ASV level. SR-MEM used a mixed-effects model with antibiotics as fixed effects, subject-level random intercepts, and autocorrelated residuals to capture temporal relationships between repeated measurements. The scale model was based on paired qPCR measurements. In contrast, Liao et al. estimated absolute abundances by multiplying relative ASV relative abundances by sample-specific qPCR totals, followed by ridge-penalized regression on time-differenced abundances. The reported antibiotic effect sizes for the Liao et al. analysis are the effects of the antibiotics over 21 days. Zero-count samples were excluded. Effect sizes are shown as heatmaps: blue indicates increased abundance associated with antibiotic use. Cells are unannotated and shown in white if either the Benjamini–Hochberg adjusted *p*-value exceeds 0.05 or the absolute effect size is below 1. The latter filter is used because Liao et al. did not report *p*-values.

Liao et al. [36] reported that intravenous vancomycin had a stronger effect on the gut microbiota than oral vancomycin—a conclusion that contradicts well-established pharmacokinetic data. Vancomycin is a water-soluble glycopeptide with poor gastrointestinal absorption; when administered intravenously, only trace amounts reach the gut [37, 38]. In contrast, oral vancomycin exerts a potent effect on the gut microbiota due to its broad-spectrum activity against gram-positive bacteria [39]. Its use as part of standard preoperative gastrointestinal decontamination further underscores its localized gut activity [40]. In line with these principles, SR-MEM yielded more biologically plausible results, indicating that oral vancomycin has a strong depleting effect on gram-positive taxa in the gut, while intravenous vancomycin has a comparatively modest impact.

Another notable inconsistency in the Liao et al. analysis concerns the carbapenem class of antibiotics. Carbapenems are potent, broad-spectrum antibiotics known to significantly deplete gut microbial diversity [38, 41]. Yet, their model identified only a single association: a modest increase in the abundance of *Lachnospiraceae*, an obligate anaerobic family well within the known spectrum of carbapenem activity [38, 42]. In contrast, SR-MEM found that carbapenem exposure was associated with broad depletion of gut microbes, including *Lachnospiraceae*, consistent with prior literature and pharmacological expectations.

Together, these results suggest SR-MEM improves inference in longitudinal studies. Unlike the original analysis, SR-MEM simultaneously addresses sparsity, counting-uncertainty, time-dependent correlation, and scale measurement uncertainty.

## 3 Discussion

In this article, we introduced *Scale Reliant Mixed-Effects Models (SR-MEM)*. SR-MEM provides a robust approach to estimating fixed effects in linear models while accounting for repeated measures that arise in complex study designs such as with nested, hierarchical, or longitudinal designs. Moreover, SR-MEM integrates recent advances from Scale Reliant Inference (SRI) to account for uncertainty in scale, thereby mitigating the problems of traditional normalizations. Through both simulated and real data analyses, we showed that SR-MEM consistently reduced false discovery rates compared to normalization-based and bias-correction-based methods while simultaneously maintaining or even improving statistical power. Across multiple analyses of previously-published data, we showed that SR-MEM produces more plausible and reproducible results. We have made SR-MEM available through the `ALDEx3` R software package.

The SR-MEM approach with scale models contributes more broadly to advancing rigor and reproducibility in statistical modeling for scientific research [43]. Similar to the principles of veridical data science, scale models and sensitivity analyses in SR-MEM address data perturbations and model stability, respectively [44, 45]. Conceptually, SR-MEM is a type of Bayesian partially identified model (PIM) [3]. By explicitly accounting for uncertainty in key parameters, Bayesian PIMs facilitate inference under weaker assumptions than fully identified models and have found use across diverse fields, including econometrics, political science, and public health [46, 47, 48, 49].

While SR-MEM provides a robust and practical framework for estimating fixed effects in the presence of repeated measures and scale uncertainty, future refinements could further enhance its utility and scope. We highlight two promising directions for future work. First, although SR-MEM is designed to account for complex experimental structures using random effects, our current formulation focuses exclusively on inference for fixed effects. Inference on random effects remains an open challenge. This is due to a subtle but important feature of the scale models used in this work: by design, these models become more conservative as the variance of the scale distribution increases. That is, as uncertainty in the scale (e.g., due to imprecision in flow cytometry measurements) increases, uncertainty in the estimated fixed effects increases monotonically. This desirable property stems from the linear structure of fixed effects within mixed-effects models. However, this monotonicity may not hold for random effects. In such settings, increasing uncertainty in scale could paradoxically reduce uncertainty in the random effects or even induce bias. These complexities suggest that our current scale models may not be appropriate for inference on random effects. Future work should investigate the conditions under which reliable inference is possible and develop specialized scale models tailored to random effects estimation.

A second promising, though less explored, direction involves leveraging recent work on predicting microbial load directly from sequence count data. For instance, Nishijima et al. [50] developed a machine learning model that predicts microbial load using features derived from sequence counts. While promising, their model achieved only modest performance, with out-of-sample *R*^2^ values in the range of 0.3–0.4, even within similar study populations. Nonetheless, such predictive models could serve as the basis for novel scale models. Rather than treating predicted microbial load values as ground truth, they could be incorporated probabilistically, wrapped in a scale model to acknowledge their uncertainty. This approach would provide a principled way to integrate machine learning predictions into statistical analyses, without requiring the unrealistic assumption that these predictions are error-free.

## 4 Methods

### 4.1 SR-MEM

#### 4.1.1 The Target Estimand and its Estimator

In SR-MEM the goal is to infer the fixed effects term *θ*_*d*_ as in Eq. (2). Core to the SRI approach is the *target estimand*: the quantity dependent on the absolute abundances *W* we wish to estimate. For SR-MEM the target estimand is explicitly defined as the generalized least squares estimator of the fixed effects *θ*_*d*_:

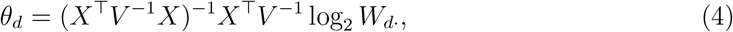

where *V* = *ZGZ*^*⊤*^ + *R*_*ρ*_ is estimated using the restricted maximum likelihood (REML) estimator to reduce bias [51].

#### 4.1.2 The SR-MEM Algorithm

The SR-MEM algorithm produces *S* independent Monte Carlo samples (referred to as “realizations” in the main text). For each Monte Carlo sample, the following steps are performed:

1. **Estimate composition:** For each sample *n ∈ {*1, …, *N }*, SR-MEM draws a posterior sample of the composition 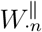 from a multinomial–Dirichlet distribution fit to the observed sequence counts *Y*:

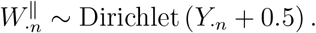
2. **Sample scale:** The total scale 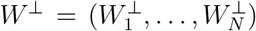 is drawn from a user-defined scale model *P*:

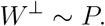
3. **Construct absolute abundances:** The sampled compositions and scales are combined using Eq. (1) to generate an estimate of the absolute abundances *W*.
4. **Fit mixed-effects model:** A mixed-effects model (Eq. (2)) is fit to W using either the `nlme` or `lme4 R` package interfaces.

Across all Monte Carlo samples, SR-MEM summarizes the fixed effects for each taxon or gene *d* by reporting the arithmetic mean of the *S* posterior estimates of *θ*_*d*_. Multiple hypothesis-corrected *p*-values are computed following the procedure described by Nixon et al. [2].

#### 4.1.3 Scale Models in SR-MEM

SR-MEM requires estimates of the *N* -length vector of scales, *W* ^*⊥*^. Any scale model *P* specified over the scale parameters *θ*^*⊥*^ (i.e., *θ*^*⊥*^ *∼ P*) can be expressed equivalently as a model over *W* ^*⊥*^ via the following relationship for each sample *n*:

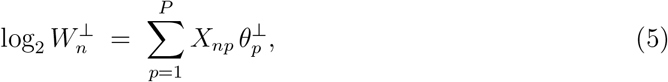

where *X*_*np*_ is the value of the *p*-th covariate for sample *n*. The resultant vector log_2_*W* ^*⊥*^ is then transformed to *W* ^*⊥*^ and used to compute the absolute abundances *W* (Eq. (1)), enabling inference within the SR-MEM framework. Further details are provided in Supplementary File 1.

### 4.2 Simulation Benchmarking

#### 4.2.1 Simulation Details

All simulations were performed using SparseDOSSA2 (v0.99.2), which generates ground-truth absolute abundances and sequence count data [26]. Data were simulated with equal-sized control and treatment groups. Each participant contributed 20 replicate samples and was assigned a random intercept drawn from *𝒩* (0, 1). For the comparison to ALDEx3 with linear modeling, intercepts were instead drawn from *𝒩* (0, 2), and all intercepts were applied using SparseDOSSA2’s spike-in feature.

We simulated three experimental scenarios. For all scenarios, SparseDOSSA2 models were trained using a correlation penalization parameter *λ* = 0.3, allowing for inter-microbe correlation. For the oral microbiome scenarios, a model was trained on 100 healthy human buccal mucosa 16S rRNA-seq samples and 197 genera, pre-processed as previously described [27]. For the skin microbiome scenario, the model was trained on 33 healthy human armpit samples and 40 families, also pre-processed as described [28]. In both cases, only taxa with no more than 70% sparsity were included for model training, and only samples with sequencing depth *≥* 1,000 reads were included.

For each scenario and each study size (ranging from 6 to 300 participants), we ran 20 independent simulations. Within each simulation, treatment effects and differentially abundant (DA) taxa were randomly assigned. Fixed treatment effects were applied using Sparse-DOSSA2’s spike-in feature, while prevalence effects were set to one-quarter of the fixed effect size to reflect expected changes in sparsity associated with positive or negative fixed effects.

For the oral microbiome scenario with 50% DA taxa, fixed effects were drawn from Uniform(0, 3) for 40% of taxa and from Uniform(*−*2, 0) for 10%. In the oral scenario with 80% DA taxa, 70% were simulated from Uniform(0, 3) and 10% from Uniform(*−*2, 0). For the skin microbiome scenario, fixed effects were drawn from Uniform(0, 3) for 50% of taxa.

To compute the true *θ*^*⊥*^ for each simulation, ground-truth absolute abundances *W* were first transformed to *W* ^*⊥*^ using Eq. (1). A mixed-effects model was then fit to log_2_ *W* ^*⊥*^, including a fixed effect for treatment and a random intercept for participants. Simulations were constrained such that *θ*^*⊥*^ *∈* [1, 2]. All samples were generated with a fixed sequencing depth of 80,000 reads.

#### 4.2.2 Analysis Details

Before model fitting, taxa with greater than 70% sparsity were amalgamated into a single “other” group. All methods, except NBZIMM and ALDEx3 with linear modeling, used the same mixed-effects model structure, which included a fixed effect for treatment and a random intercept for each participant. NBZIMM additionally included a fixed-effect offset for log_2_ sequencing depth, as recommended [11]. ALDEx3 with linear modeling did not include a random intercept.

Both SR-MEM and ALDEx3 assumed the scale model *θ*^*⊥*^ *∼ 𝒩* (1.3, 0.5), which was transformed to a corresponding scale model over *W* ^*⊥*^ using Eq. (5). SR-MEM and ALDEx3 with linear modeling were implemented using ALDEx3 (v0.4.0) with 300 Monte Carlo samples. SR-MEM used the `lme4` package for model fitting.

ANCOM-BC2 (v2.6.0) was run with default parameters; taxa that failed the pseudo-count sensitivity test were assigned *p*-values of 1 after multiple hypothesis test correction [14]. MaAsLin2 (v1.18.0) was also run with default settings, including total sum scaling (TSS) normalization. NBZIMM (v1.0) used the `mms` function with a zero-inflated negative binomial model. LinDA (v1.2) was run with zero imputation enabled. lmerSeq (v0.1.7) applied DESeq2 size-factor normalization followed by variance-stabilizing transformation (VST), as recommended [13, 52].

For all methods, taxa grouped as “other” were assigned *p*-values of 1 following multiple testing correction. A taxon was considered a true positive if the null hypothesis of no treatment effect was rejected at FDR *≤* 0.05 and the sign of the estimated treatment effect matched the ground truth. If the sign was incorrect, the taxon was included in the denominator for power calculations but excluded from the numerator. All methods used the Benjamini-Hochberg procedure for multiple testing correction.

### 4.3 Reanalysis of Oral Microbiome Perturbation Study

Sequence count data were pre-processed following Marotz et al. [31], and analyses were conducted at the genus level. Taxa with more than 70% sparsity were grouped into a single “other” category, resulting in a total of 59 taxa. The dataset comprised 81 samples from 21 participants. For each sample, oral microbial load was measured in duplicate using flow cytometry. Denoting 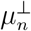 as the mean and 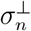 as the standard deviation of the two log-transformed replicate cell counts for sample *n*, we modeled the absolute abundance as:

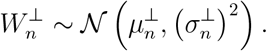

All methods except NBZIMM used the same mixed-effects model structure, including fixed effects for time point, perturbation, and their interaction, as well as a random intercept for each participant. The fixed effects intercept corresponded to the baseline (pre-treatment) time point under the reference perturbation (water). NBZIMM additionally included a fixed-effect offset for log_2_ sequencing depth.

For SR-MEM, 2,000 Monte Carlo samples were drawn using the `lme4`-based implementation in ALDEx3 (v0.4.0). For the QMP analysis, absolute abundance estimates *W* were estimated using the QMP method of Vandeputte et al. [16], based on the average of the two microbial load replicates per sample. A pseudo-count of 0.5 was added prior to log-transformation. The same mixed-effects model described above was applied to each taxon.

NBZIMM (v1.0) used the `mms` function with a zero-inflated negative binomial model. ANCOM-BC2 (v2.2.1) was run with default settings; taxa failing the pseudo-count sensitivity test were assigned a *p*-value of 1 following multiple testing correction. LinDA (v1.2) applied zero imputation. lmerSeq (v0.1.7) used DESeq2 size-factor normalization followed by variance-stabilizing transformation (VST), as recommended [13, 52]. All methods applied multiple testing correction using the Benjamini–Hochberg procedure.

### 4.4 Reanalysis of IBDMDB Data

#### 4.4.1 Metagenomic Data Pre-processing

Raw metagenomic sequence data were obtained from the IBDMDB database [35]. Host DNA was removed using Kneaddata (v0.12.0), and taxonomic profiling with read count estimation was performed using MetaPhlAn (v4.0.6) and the CHOCOPhlAn database (v30) [53, 54]. Genera with zero counts in more than 75% of samples were grouped into a single “other” category, resulting in 43 taxa analyzed across 1,297 samples from 106 individuals. Sample metadata, including gut dysbiosis classification (dysbiotic vs. non-dysbiotic), were also obtained from the IBDMDB database. Analyses in the main text compared 582 Crohn’s disease (CD) samples to 362 non-IBD samples; the remaining 353 samples from ulcerative colitis (UC) patients are reported separately (Supplementary File 3).

#### 4.4.2 Analyses of IBDMDB Data

Disease diagnosis (non-IBD, UC, or CD) and dysbiosis status were combined into a single covariate labeled *disease-dysbiosis*. All analyses used a mixed-effects model including fixed effects for antibiotic use and disease-dysbiosis, and random intercepts for participant and site. The fixed-effects intercept corresponded to non-IBD samples collected during non-dysbiosis without antibiotic use.

SR-MEM used 1,000 Monte Carlo samples and a prior scale model informed by visual inspection of Figure 2 in Sarrabayrouse et al. [34]:

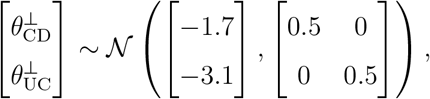

where 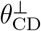and 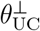 represent the difference in average log_2_scale for CD and UC samples during dysbiosis, relative to non-IBD samples during non-dysbiosis. CD and UC samples taken during non-dysbiosis and non-IBD samples taken during dysbiosis were assumed to have differences in average log_2_ scale of zero. This scale model was transformed to one over *W* ^*⊥*^ using Eq. (5).

In the ASR-MEM analysis, observed counts were converted to relative abundances 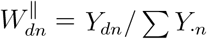, then transformed using the arcsine square-root (ASR) transformation:

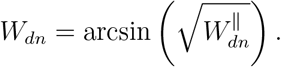

These transformed values were used in the same mixed-effects model as SR-MEM.

The SR-MEM-CLR analysis was identical to SR-MEM, except that absolute abundances were estimated using centered log-ratio (CLR) normalization:

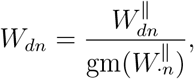

where gm denotes the geometric mean across taxa within each sample.

Multiple hypothesis testing correction for all analyses was performed using the Benjamini-Hochberg procedure.

#### 4.4.3 Reanalysis of Vandeputte Data

Sequence count data were pre-processed as described by Vandeputte et al. [16]. Genera with zero counts in more than 75% of samples were aggregated into a single “other” category, resulting in 30 taxa analyzed across 95 samples. The scale model, previously defined by Nixon et al. [2], was:

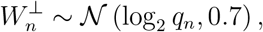

where *q*_*n*_ denotes the flow cytometry measurement of cells per gram of frozen feces for sample *n*. ALDEx2 (v1.40.0) was used to compare disease status (CD vs. control) with 1,000 Monte Carlo samples. Multiple testing correction was performed using the Benjamini-Hochberg procedure.

### 4.5 Reanalysis of Longitudinal Antibiotic Dataset

#### 4.5.1 Data Pre-processing

16S rRNA sequencing count data were pre-processed by Liao et al. and obtained from Figshare (version 6) [36]. Taxa were analyzed at the amplicon sequence variant (ASV) level. Antibiotics were categorized by route of administration: oral (glycopeptides, penicillins, quinolones, sulfonamides, and macrolide derivatives) or intravenous (aztreonam, carbapenems, cephalosporins, glycopeptides, metronidazole, oxazolidinones, penicillins, and quinolones). All other antibiotics were grouped into an “other” category. Antiviral and antifungal treatments were excluded from filtering and modeling.

#### 4.5.2 SR-MEM Analysis

Samples lacking paired qPCR measurements were excluded. A sample was labeled as unaffected by antibiotics if it was collected either before antibiotic administration or at least 21 days afterward. Samples collected within 21 days post-antibiotic-treatment were excluded. Patients with fewer than 10 samples were also removed. After filtering, the final dataset included 2,589 samples from 144 patients.

The SR-MEM model included 14 fixed effects: one for each antibiotic-route combination and one for the “other” antibiotic category. A random intercept was included for each patient. SR-MEM analysis was performed using the `nlme`-based implementation of ALDEx3 (v0.4.0). A first-order autoregressive correlation structure with the formula *∼* TimePoint|PatientID was used. This defines the residual covariance matrix as *R*_*ρ*_ = *σ*^2^ *L*_*ρ*_, where each element *l*_*xy*_ of *L*_*ρ*_ is:

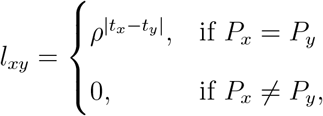

with *t*_*x*_ and *t*_*y*_ representing the collection time points (in days), and *P*_*x*_ and *P*_*y*_ denoting the patients corresponding to samples *x* and *y*, respectively. The autocorrelation parameter *ρ* was estimated via restricted maximum likelihood (REML). We used 3,000 Monte Carlo samples in the SR-MEM analysis. For each sample *n* with an observed qPCR value *q*_*n*_, the scale model was:

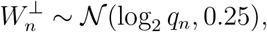

where *q*_*n*_ is the qPCR-measured microbial load (cells per gram of stool). Multiple testing correction was performed using the Benjamini-Hochberg procedure.

#### 4.5.3 Liao Analysis

The original Liao et al. analysis was reproduced as described in their publication, using the provided MATLAB code [36]. Multiple testing correction was performed using the Benjamini-Hochberg procedure.

## 5 Data Availability

All datasets used in this study are publicly available and have been previously published. The oral microbiome data used to train SparseDOSSA2 are available from Qiita (study ID 10370), and the skin microbiome training data are accessible via the `HMP2Data R` package. Data from the oral microbiome perturbation study are available from Qiita (study ID 11899). The IBDMDB dataset is available at https://ibdmdb.org. Data from the longitudinal antibiotic study are available on Figshare at DOI: 10.6084/m9.figshare.c.5271128.v6.

## 6 Code Availability

All code used to generate figures, supplementary figures, and supplementary files is available at: https://github.com/Silverman-Lab/SR-MEM. The implementation of SR-MEM is included in the `ALDEx3 R` package, available at: https://github.com/jsilve24/ALDEx3.

## 7 Contributions

KCM wrote all code and performed all analyses in the manuscript. JDS obtained funding. KCM and JDS contributed the idea for the scale-reliant mixed effects modeling statistical framework. All authors read and approved the manuscript.

## 8 Ethical Declarations

The authors declare that they have no conflict of interest.

## 10 Acknowledgments

The authors would like to thank Dr. Rachel Silverman for her manuscript comments.

