## Supplementary File 1 for "Scale Reliant Mixed Effects Models Enhance Microbiome Data Analysis"

### 1 Designing Scale Models for SR-MEM

Defining a scale model for $W^{\perp} = (W_1^{\perp}, \ldots, W_N^{\perp})$ may seem daunting, as it requires specifying a joint distribution over $N$ unobserved scale parameters. However the field of SRI has developed practical strategies that can simplify this task considerably.

For example, suppose external measurements of scale, denoted $q_n$, are available—such as from flow cytometry or qPCR. Further, assume that replication experiments have been conducted to estimate the measurement error, and that the variance of the log-transformed measurements is known, i.e., $\text{Var}(\log_2 q_n) = \sigma^2$. In this setting, it is often reasonable to assume conditional independence and define the scale model as $\log_2 W_n^{\perp} \sim \mathcal{N}(\log_2 q_n, \sigma^2)$. This strategy has been successfully employed in multiple SRI studies [1, 2, 3].

When external measurements of scale are unavailable and scale models must instead be specified using prior knowledge or biologically plausible assumptions, we generally recommend two strategies.

First, scale models can be constructed to generalize a desired normalization procedure. As discussed in prior work [1, 3, 4], many common normalizations can be expressed as transformations of the form:

$$W_{\cdot n} = \frac{W^{\parallel}_{\cdot n}}{\phi(W^{\parallel}_{\cdot n})}, \tag{1}$$

for some scalar-valued function $\phi$. For example, Total Sum Scaling (TSS) normalization corresponds to the case where $\phi(W^{\parallel}_{\cdot n}) = 1$, while Centered Log-Ratio (CLR) normalization is obtained when

$$\phi(W^{\parallel}_{\cdot n}) = \left(\prod_{d=1}^{D} W^{\parallel}_{dn}\right)^{1/D},$$

i.e., the geometric mean of the components of $W^{\parallel}_{\cdot n}$. Normalizations based on housekeeping genes also fit within this framework. Let $H \subset \{1, \ldots, D\}$ denote a set of taxa (or genes) believed to be invariant with respect to the covariates in $X$. In this case, a natural choice of scaling function is

$$\phi(W^{\parallel}_{\cdot n}) = \left(\prod_{d \in H} W^{\parallel}_{dn}\right)^{1/|H|},$$

which assumes that the geometric mean abundance of the housekeeping set remains stable across experimental conditions. Regardless of the specific normalization used, Eq. (1) is unlikely to hold exactly. To account for possible deviations, we recommend generalizing the normalization by specifying a scale model of the form

$$\log_2 W^{\perp}_{n} \sim \mathcal{N}(-\log_2 \phi(W^{\parallel}_{\cdot n}), \sigma^2),$$

where $\sigma^2$ is a user-defined variance parameter representing the degree of uncertainty in the normalization. This formulation encodes prior belief in the normalization procedure while allowing for plausible departures from it. Additional examples and discussion of this approach can be found in prior work [1, 2, 3].

Second, scale models can be constructed using a reparameterization trick that simplifies the problem. Rather than directly specifying a distribution over the $N$-dimensional vector $W^{\perp} = (W^{\perp}_1, \ldots, W^{\perp}_N)$, we instead focus on how $W^{\perp}$ contributes to estimation of the fixed effects $\theta_d$. Specifically, the standard estimator for fixed effects in mixed-effects models (e.g., the maximum likelihood estimator) can be expressed as a linear transformation: $\theta_d = A \log W_{d\cdot}$, where $A = (X^{\top} V^{-1} X)^{-1} X^{\top} V^{-1}$ and $V = ZGZ^{\top} + R_{\rho}$. By linearity, this estimator decomposes as

$$\begin{aligned}\theta_d &= A \log_2 W_{d\cdot} \\ &= \underbrace{A \log_2 W^{\parallel}_{d\cdot}}_{\theta^{\parallel}_d} + \underbrace{A \log_2 W^{\perp}}_{\theta^{\perp}}, \end{aligned} \tag{2}$$

where $\theta^{\parallel}_d$ and $\theta^{\perp}$ are each $P$-dimensional vectors representing the compositional and scale components of the fixed effects, respectively. That is, $\theta^{\parallel}_d$ and $\theta^{\perp}$ describe how $\log_2 W^{\parallel}_{dn}$ and $\log_2 W^{\perp}_n$, respectively, vary with the covariates in $X$.

Instead of modeling $W^{\perp}$ directly, we can instead model $\theta^{\perp}$ which only $P$-dimensional. To guide model specification, we recommend thinking of $\theta^{\perp}$ as the coefficients in a regression model of the form:

$$\log_2 W^{\perp}_n = \sum_{p=1}^{P} X_{np} \times \theta^{\perp}_p \tag{3}$$

where each element $\theta^{\perp}_p$ represents the effect of covariate $X_{np}$ on the log scale. Once a distribution $\theta^{\perp} \sim P$ is specified, realizations of $W^{\perp}$ can be generated by first sampling $\theta^{\perp}$ from $P$, then applying Eq. (3) to compute $\log_2 W^{\perp}_n$ for all $n$. Note Eq. (3) is the same as Eq. (5) in *Methods* in the main text.

**Example.** Consider differential abundance analysis which can be framed as a linear model with covariates $X_n = (1, c_n)$, where $c_n \in \{0, 1\}$ denotes treatment condition (e.g., placebo versus drug). In this case, Eq. (3) becomes:

$$\log_2 W^{\perp}_n = \theta^{\perp}_1 + \theta^{\perp}_2 c_n,$$

where $\theta^{\perp}_2$ is of primary scientific interest—it quantifies the difference in scale (e.g., total microbial load) between treatment groups. In a motivating example from [2], the authors studied changes in bacterial load as an inoculated microbial community expanded to occupy a larger volume. Based on the ratio of inoculation volume to final vessel volume, they specified the prior $\theta^{\perp}_2 \sim \mathcal{N}\left(\log_2\left(\frac{100 \text{ mL}}{400 \text{ mL}}\right), 0.5^2\right)$, where the variance accounts for possible deviations due to unmeasured differences in carrying capacity across environments. Since the intercept $\theta_1^\perp$ is not of scientific interest, it can be fixed to zero without loss of generality. To sample from the corresponding scale model: draw $\theta_2^\perp \sim \mathcal{N}(\log_2 0.25, 0.5^2)$, and compute $\log_2 W_n^\perp = \theta_2^\perp c_n$, for each $n = 1, \ldots, N$. This procedure yields realizations of $W^\perp$ consistent with prior knowledge, and allows SR-MEM to propagate this uncertainty through to inference on $\theta_d$.
