## Supplementary figures and images for "Scale Reliant Mixed Effects Models Enhance Microbiome Data Analysis"

### Supplementary Figure 1

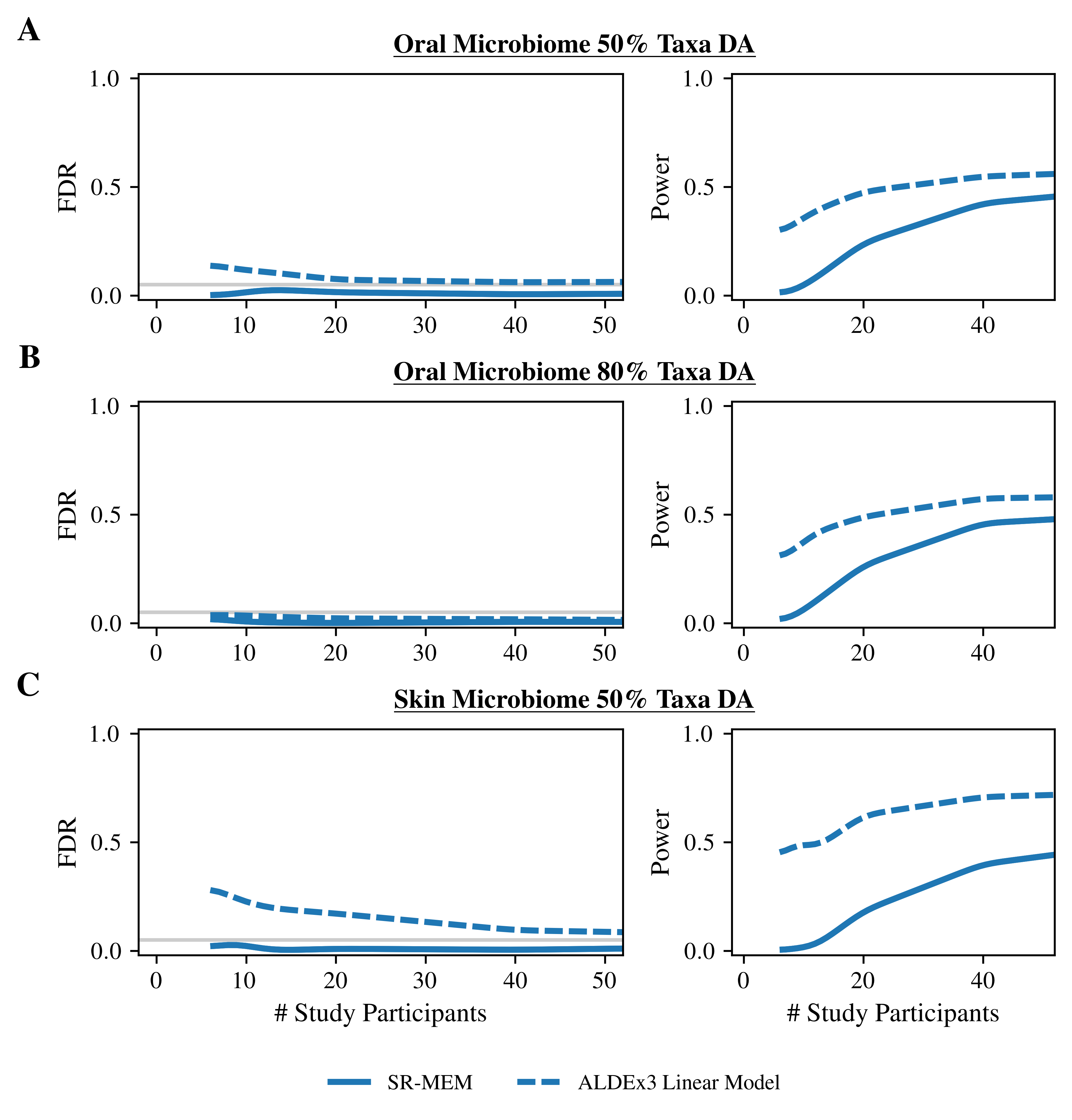

### Supplementary Figure 2

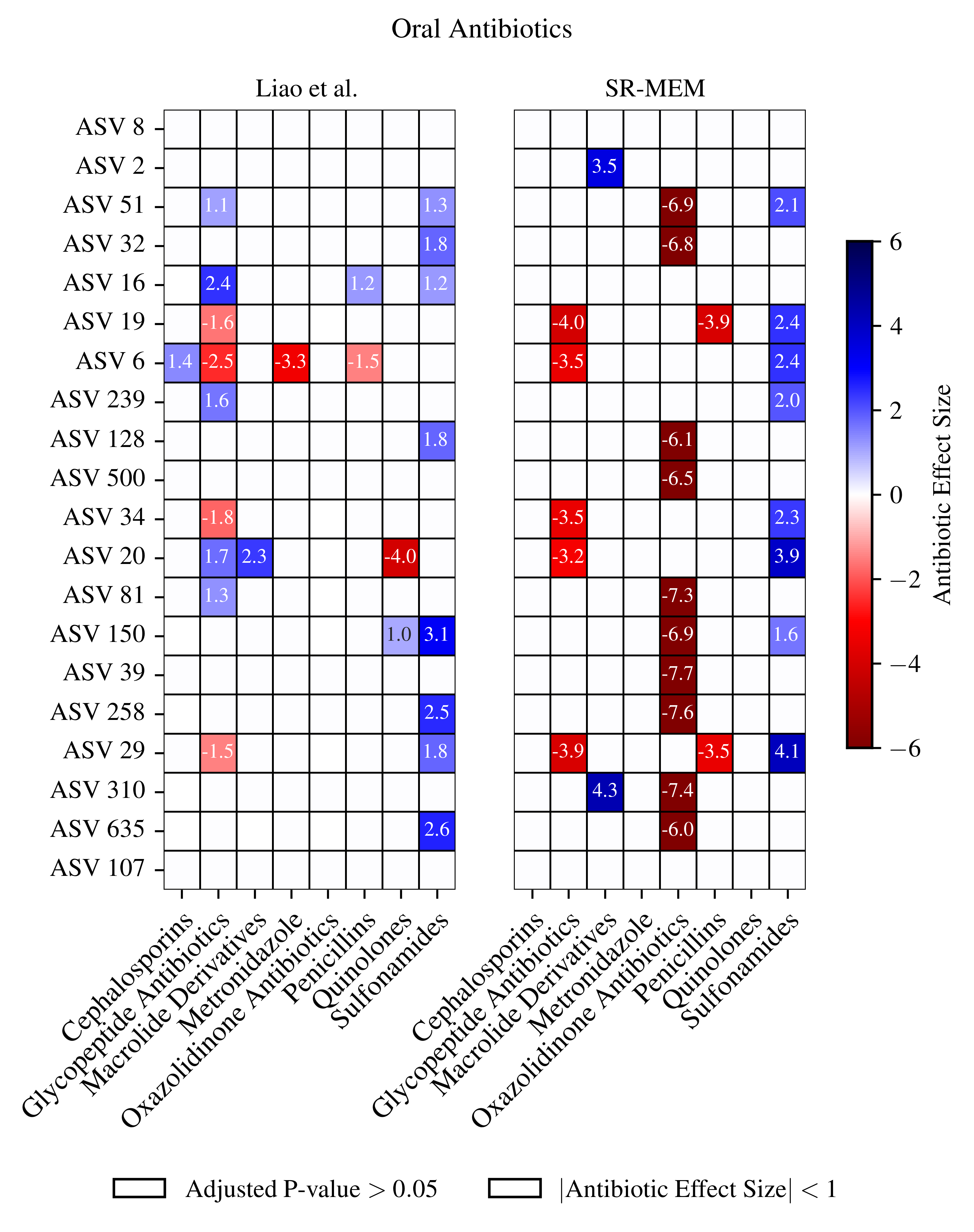

### Supplementary Figure 3

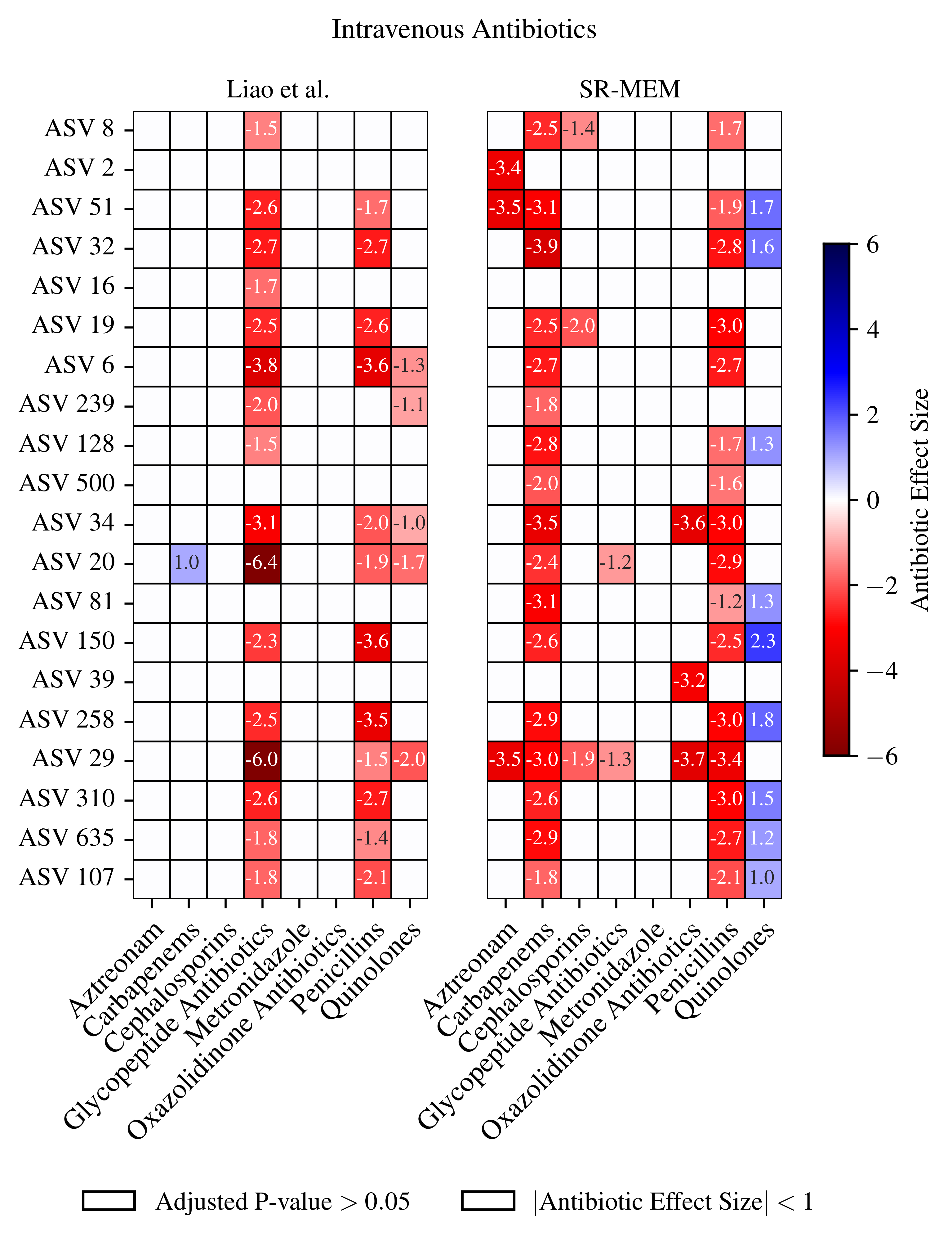
